# Curation of BIDS (CuBIDS): a workflow and software package for streamlining reproducible curation of large BIDS datasets

**DOI:** 10.1101/2022.05.04.490620

**Authors:** Sydney Covitz, Tinashe M. Tapera, Azeez Adebimpe, Aaron Alexander-Bloch, Maxwell A. Bertolero, Eric Feczko, Alexandre R. Franco, Raquel E. Gur, Ruben C. Gur, Timothy Hendrickson, Audrey Houghton, Kahini Mehta, Kristin Murtha, Anders J. Perrone, Tim Robert-Fitzgerald, Jenna M. Schabdach, Russell T Shinohara, Jacob W. Vogel, Chenying Zhao, Damien A. Fair, Michael P. Milham, Matthew Cieslak, Theodore D. Satterthwaite

## Abstract

The Brain Imaging Data Structure (BIDS) is a specification accompanied by a software ecosystem that was designed to create reproducible and automated workflows for processing neuroimaging data. BIDS Apps flexibly build workflows based on the metadata detected in a dataset. However, even BIDS valid metadata can include incorrect values or omissions that result in inconsistent processing across sessions. Additionally, in large-scale, heterogeneous neuroimaging datasets, hidden variability in metadata is difficult to detect and classify. To address these challenges, we created a Python-based software package titled “Curation of BIDS” (CuBIDS), which provides an intuitive workflow that helps users validate and manage the curation of their neuroimaging datasets. CuBIDS includes a robust implementation of BIDS validation that scales to large samples and incorporates DataLad––a version control software package for data––to ensure reproducibility and provenance tracking throughout the entire curation process. CuBIDS provides tools to help users perform quality control on their images’ metadata and identify unique combinations of imaging parameters. Users can then execute BIDS Apps on a subset of participants that represent the full range of acquisition parameters that are present, accelerating pipeline testing on large datasets.

**HIGHLIGHTS:**

- CuBIDS is a workflow and software package for curating BIDS data.
- CuBIDS summarizes the heterogeneity in a BIDS dataset.
- CuBIDS prepares BIDS data for successful preprocessing pipeline runs.
- CuBIDS helps users perform metadata-based quality control.

## INTRODUCTION

The Brain Imaging Data Structure (BIDS) specification provides a standardized format for organizing and describing neuroimaging data (Gorgolewski et al., 2016). BIDS relies on specific nested directory structures and filename conventions and requires that each Neuroimaging Informatics Technology Initiative (NIfTI) file be accompanied by a JavaScript Object Notation (JSON) sidecar––a data dictionary detailing its corresponding image’ s metadata. BIDS is especially helpful when dealing with large, multimodal studies; as the number of subjects and scans increases, generalizable structures and standards become not only beneficial but essential. Pipelines that ingest BIDS datasets––commonly referred to in the BIDS software ecosystem as “BIDS Apps”––such as fMRIPrep and QSIPrep––rely heavily on correct specification of BIDS, as they build workflows based on the metadata encountered (Esteban et al., 2019; Cieslak et al., 2021). While generally an important and useful feature, this workflow construction structure can also be a vulnerability: if the BIDS metadata is inaccurate, a BIDS app may build an inappropriate (but technically “correct”) preprocessing pipeline. As a result, users must understand and verify that the metadata present in BIDS is correct. This usually requires meticulous curation––the process of checking and fixing filename or metadata issues present in a dataset.

While large, multi-modal neuroimaging datasets constitute extremely valuable data resources, they also frequently possess substantial heterogeneity in their image acquisition parameters. BIDS provides an ideal structure for organizing neuroimaging data, but the size and complexity of large-scale datasets can render curation both tedious and difficult. Data curation can be an ad-hoc process that involves substantial manual intervention; such manual curation is usually neither well tracked nor reproducible. Thus, curation constitutes a major vulnerability in the field-wide effort to create fully reproducible analytic workflows for neuroimaging data. Finally, many current BIDS tools (e.g. the BIDS Validator and PyBIDS) that were created for the curation process were optimized for small fMRI studies and may behave erratically when given large quantities of heterogeneous data (Yarkoni et al., 2019).

With these challenges in mind, we developed “Curation of BIDS” (CuBIDS): a software package that provides sanity preserving workflows to help users curate large BIDS datasets. CuBIDS provides users with customizable features to visualize heterogeneity in complex BIDS datasets and includes a robust, scalable implementation of BIDS validation that can be applied to arbitrarily-sized datasets. Critically, CuBIDS renders curation reproducible via an easy-to-use, wrapped implementation of DataLad (Halchenko et al., 2021). Finally, CuBIDS provides tools to identify unique combinations of imaging parameters in a dataset so that users can test BIDS Apps on a subset of participants that represents the parameter space of the entire dataset. This option dramatically speeds up pipeline testing, as users can be assured that they have tested a BIDS App on the full range of acquisition parameters present in a dataset. As described below, CuBIDS facilitates an understanding of what is present in a BIDS dataset, allows for reproducible BIDS curation, and accelerates successful data processing at scale.

## MATERIALS AND METHODS

The standard lifecycle of a neuroimaging study begins with acquisition and ends with image analysis and hypothesis testing. CuBIDS’ role in this process begins directly after the data has been organized into a BIDS directory structure with BIDS-like filenames. The CuBIDS workflow ends when the entire dataset is ready to be run through modality-specific preprocessing pipelines. As curation occurs quite early in this timeline of preparing neuroimaging data for analysis, decisions made during curation will affect every subsequent stage.

### Data and Code Availability Statement

A copy of the small, example dataset whose curation we walk through in the Results section is compressed into a ZipFile and submitted with this paper under “Supplementary Material.” Additionally, the Philadelphia Neurodevelopmental Cohort (PNC), the dataset whose curation we summarize in the second portion of the Results, is publicly available in the Database of Genotypes and Phenotypes (dbGaP accession phs000607.v3.p2). The source code for CuBIDS is publicly available at https://github.com/PennLINC/CuBIDS, the documentation for our software is available at https://cubids.readthedocs.io/en/latest/, and our package is available for download on the Python Package Manager (pypi) https://pypi.org/project/cubids/.

### Ethics Statement

No new data was collected specifically for this paper. The Philadelphia Neurodevelopmental Cohort (PNC) (Satterthwaite et al., 2014) was approved by IRBs of The University of Pennsylvania and Children’ s Hospital of Philadelphia. All adult participants in the PNC provided informed consent to participate; minors provided assent alongside the informed consent of their parents or guardian.

### Overview

CuBIDS provides a workflow that aids users in curating large, heterogeneous BIDS datasets. CuBIDS summarizes a dataset’ s metadata, enabling users to visualize and understand the variability in critical scanning parameters and fix errors when they are present. To do this,

CuBIDS features several command line interface (CLI) programs (**Table. 1**). Notably, all CuBIDS CLI programs wrap DataLad as an optional dependency so that the user can implement reproducible tracking at any stage of curation or revert to a prior state of their data. If the user wants to apply DataLad version control while using CuBIDS, they can run the CLI programs with the `–-use-datalad` optional flag set.

**Table 1:** cubids command line interface programs. cubids features several command line interface (cli) programs that help users curate and process bids datasets. we use the color-coded backgrounds to map each program to a stage in the curation workflow seen in **Fig. 1**. These programs were built for the steps of a study’ s curation process. some programs––such as `print-metadata-fields`, `group`, `validate`, and `copy-exemplars`––require only “read” access to the data and aid the user in visualizing a dataset’ s heterogeneity. others––such as `apply`, `purge`, `undo`, and `remove-metadata-fields`––require write access, as they involve modifying metadata, changing filenames, or removing entire subjects altogether.

| COMMAND | POSITIONAL ARGUMENTS | OPTIONAL ARGUMENTS | OUTPUT FILES | DESCRIPTION |
| --- | --- | --- | --- | --- |
| `cubids-print-metadata-fields` | `bids_dir`<br>the root of a BIDS dataset. It should contain sub-X directories and `dataset_description.json` |  |  | Prints out all sidecar field names present in the dataset |
| `cubids-remove-metadata-fields` | `bids_dir`<br>the root of a BIDS dataset. It should contain sub-X directories and `dataset_description.json` | `--fields`<br>`FIELDS`<br>`[FIELDS ...]`<br>space-separated list of metadata fields to remove (default: []) |  | Removes a desired list of metadata fields and can be used to remove PHI from datasets |
| cubids-add-nifti-info | bids_dir<br>the root of a BIDS dataset. It should contain sub-X directories and dataset_description.json | --use-datalad<br>ensure that there are no untracked changes before finding groups and save dataset after NifTI info is added to sidecars (default: False)<br>--force-unlock<br>unlock dataset before adding nifti info (default: False) |  | Adds the following information from the headers of images in NIfTI format: Obliquity, NumVolumes, Dim*Size, and VoxelSizeDim* to its corresponding sidecar |
| cubids-validate | bids_dir<br>the root of a BIDS dataset. It should contain sub-X directories and dataset_description.json<br>output_prefix<br>file prefix to which tabulated validator output is written | --sequential<br>run the BIDS Validator sequentially on each subject (default: False)<br>--ignore_nifti_headers<br>disregard NifTI header content during validation (default: False)<br>--ignore_subject_consistency<br>skip checking that any given file for one subject is present for all other subjects (default: False)<br>--sequential-subjects<br>SEQUENTIAL_SUBJECTS<br>[SEQUENTIAL_SUBJECTS<br>...]<br>List: Filter the sequential run to only include the listed subjects.<br>e.g. --sequential-subjects sub-01 sub-02 sub-03<br>(default: None) | output_prefix_validation.csv<br>CSV file with one row per file containing a BIDS validation error. Columns: files, type, severity, description, code, url, subject | Stable, robust version of the BIDS-validator |
| cubids-group | bids_dir<br>the root of a BIDS dataset. It should contain sub-X directories and dataset_description.json<br>output_prefix<br>file prefix to which tabulated grouping output is written | --acq-group-level<br>ACQ_GROUP_LEVEL<br>Level at which acquisition groups are created<br>options: "subject" or "session"<br>(default: subject)<br>--config<br>CONFIG<br>path to a config file for grouping<br>(default: None) | output_prefix_summary.csv<br>output_prefix_files.csv<br>output_prefix_AcqGrouping.csv<br>output_prefix_AcqGroupInfo.txt | Produces four CSVs that display the heterogeneity present in the dataset in a user-friendly format |
| cubids-purge | bids_dir<br>the root of a BIDS dataset. It should contain sub-X directories and dataset_description.json<br>scans<br>absolute path<br>to the txt file of scans whose associations should be purged | --use-datalad<br>save deletions after purge runs<br>(default: False) |  | Takes in a list of NIfTI images and removes them and all their association files and IntendedFor references from the dataset |
| cubids-apply | bids_dir<br>the root of a BIDS dataset. It should contain sub-X directories and dataset_description.json<br>edited_summary_csv<br>the *_summary.csv that has been edited in the MergeInto and RenameKeyGroup columns.<br>files_csv<br>the *_files.csv that the *_summary.csv corresponds to.<br>output_prefix<br>the file prefix for writing the post-apply grouping outputs | --use-datalad<br>save changes applied to the dataset (default: False)<br>--acq-group-level<br>ACQ_GROUP_LEVEL<br>level at which acquisition groups are created<br>options: "subject" or "session" (default: subject)<br>--config<br>CONFIG<br>path to a config file for grouping (default: None) | output_prefix_summary.csv<br>output_prefix_files.csv<br>output_prefix_AcqGrouping.csv<br>output_prefix_AcqGroupInfo.txt | Applies the user's edits to the *_summary.csv file to the BIDS dataset |
| cubids-copy-exemplars | bids_dir<br>absolute path to the root of a BIDS dataset. It should contain sub-X directories and dataset_description.json<br>exemplars_dir<br>absolute path to the root of a BIDS dataset containing one subject from each Acquisition Group. It should contain sub-X directories and dataset_description.json<br>exemplars_csv<br>absolute path to the .csv file that lists one subject from each Acquisition Group (*_AcqGrouping.csv from the cubids-group output) | --use-datalad<br>ensure that there are no untracked changes before finding groups and save exemplar dataset post creation (default: False)<br>--min-group-size<br>MIN_GROUP_SIZE<br>minimum number of subjects an Acquisition Group must have in order to be included in the exemplar dataset (default: 1) |  | copies one subject from each exemplar group into a separate directory |
| cubids-undo | bids_dir<br>absolute path<br>to the root of a<br>BIDS dataset.<br>It should<br>contain sub-X<br>directories and<br>dataset_descri<br>ption.json |  |  | reverts data-set<br>to its state prior<br>to the most<br>recent saved<br>modifications<br>(only available<br>to users using<br>the DataLad-<br>based version<br>control option) |

### Software development practices

We applied test-driven development while building CuBIDS, prioritizing writing tests for each new feature concurrent with its construction. We integrated CircleCI––a web-based continuous integration testing platform––into our GitHub repository so that each new commit is run through the full suite of tests. We apply a standardized approach to fixing bugs and adding features: first creating an issue on our GitHub page and then creating a new branch of our code base named specifically for fixing that issue. Once the issue is fixed on the new branch, a pull request merges the new branch into the main branch with the issue tagged. If all continuous integration tests pass and the merge is successful, the issue gets automatically closed. Centering our development process around both tests and issues has ensured the integrity of the code and facilitated both organization and documentation.

### Installation, setup, and version control

We recommend users install CuBIDS inside an Anaconda-based Python environment. Users can install Anaconda/Miniconda/Miniforge, create and activate an environment, and then obtain CuBIDS locally by either installing from the <u>Python Package Manager (Pypi)</u> using pip or cloning directly from the <u>CuBIDS GitHub repository</u>. Documentation regarding use of CuBIDS is publicly available on our <u>Read the Docs page</u>. Notably, CuBIDS commands incorporate version control using DataLad as an optional dependency. Operationalizing command line programs with the `–use-datalad` flag set allows users to track changes they make to their dataset and revert their dataset back to earlier versions if necessary. If users have their BIDS data checked into DataLad, which will enable version control, they can also leverage `cubids-undo`. If users run one of the CuBIDS programs that makes changes to the metadata or filenames and with the `–-use-datalad` flag, those changes will be saved as a commit. If users decide they want to undo those changes and return the dataset to its prior state, they can run `cubids-undo`, which will revert the most recent commit to the dataset. If users would like to access this functionality, they must separately install both DataLad and Git Annex (a dependency of DataLad). Although users can run CuBIDS programs without DataLad, opting to leverage the version control capabilities is recommended, as it allows for fully reproducible curation.

### Definitions

The CuBIDS workflow centers on three main concepts. The first is a “Key Group” –– the set of scans whose filenames share all BIDS filename key-value pairs, except for subject and session. For example, CuBIDS would place a T1w NIfTI file named `sub-X_ses-A_acq-refaced_T1w.nii.gz`, which contains the BIDS key-value pair “acq-refaced”––in the following Key Group: `acquisition-refaced_datatype-anat_suffix-T1w`. Notably, Key Groups only consider the scan’ s BIDS filename; they do <u>not</u> account for the variance in metadata fields that might be present in the JSON sidecars.

For this reason, within each Key Group, we define a “Parameter Group” as the set of scans with identical metadata parameters contained in their sidecars. Parameter Groups exist within Key Groups and are denoted numerically––each Key Group will have *n* Parameter Groups, where *n* is the number of unique sets of scanning parameters present in that Key Group. For example, a T1w can belong to Key Group `acquisition-refaced_datatype-anat_suffix-T1w` and Parameter Group 1. CuBIDS defines Parameter Groups within Key Groups because differences in parameters can affect how BIDS Apps will configure their pipelines (e.g. Fieldmap availability, multiband factor, etc).

Next, we define a “Dominant Group” as the Parameter Group that contains the most scans in its Key Group. Analogously, we define a “Variant Group” as any Parameter Group that is non-dominant. This is an important term because (as described below) CuBIDS can optionally rename all Variant Groups in an automated and reproducible fashion.

Finally, we define an “Acquisition Group” as a collection of sessions across participants that contain the exact same set of Key and Parameter Groups. Since Key Groups are based on the BIDS filenames—and therefore both modality and acquisition specific—each BIDS session directory contains images that belong to a set of Parameter Groups. CuBIDS assigns each session––or set of Parameter Groups––to an Acquisition Group such that all sessions in an Acquisition Group possesses an identical set of scan acquisitions and metadata parameters across <u>all</u> image modalities present in the dataset. We find Acquisition Groups to be a particularly useful categorization of BIDS data, as they identify homogeneous sets of sessions (not individual scans) in a large dataset. They are also useful for expediting the testing of pipelines; if a BIDS App runs successfully on a single subject from each Acquisition Group, one can be confident that it will handle all combinations of scanning parameters in the entire dataset. These various sets of methods by which one can group a BIDS dataset are critical to the CuBIDS workflow (see **Fig 1**).

**Fig. 1.**
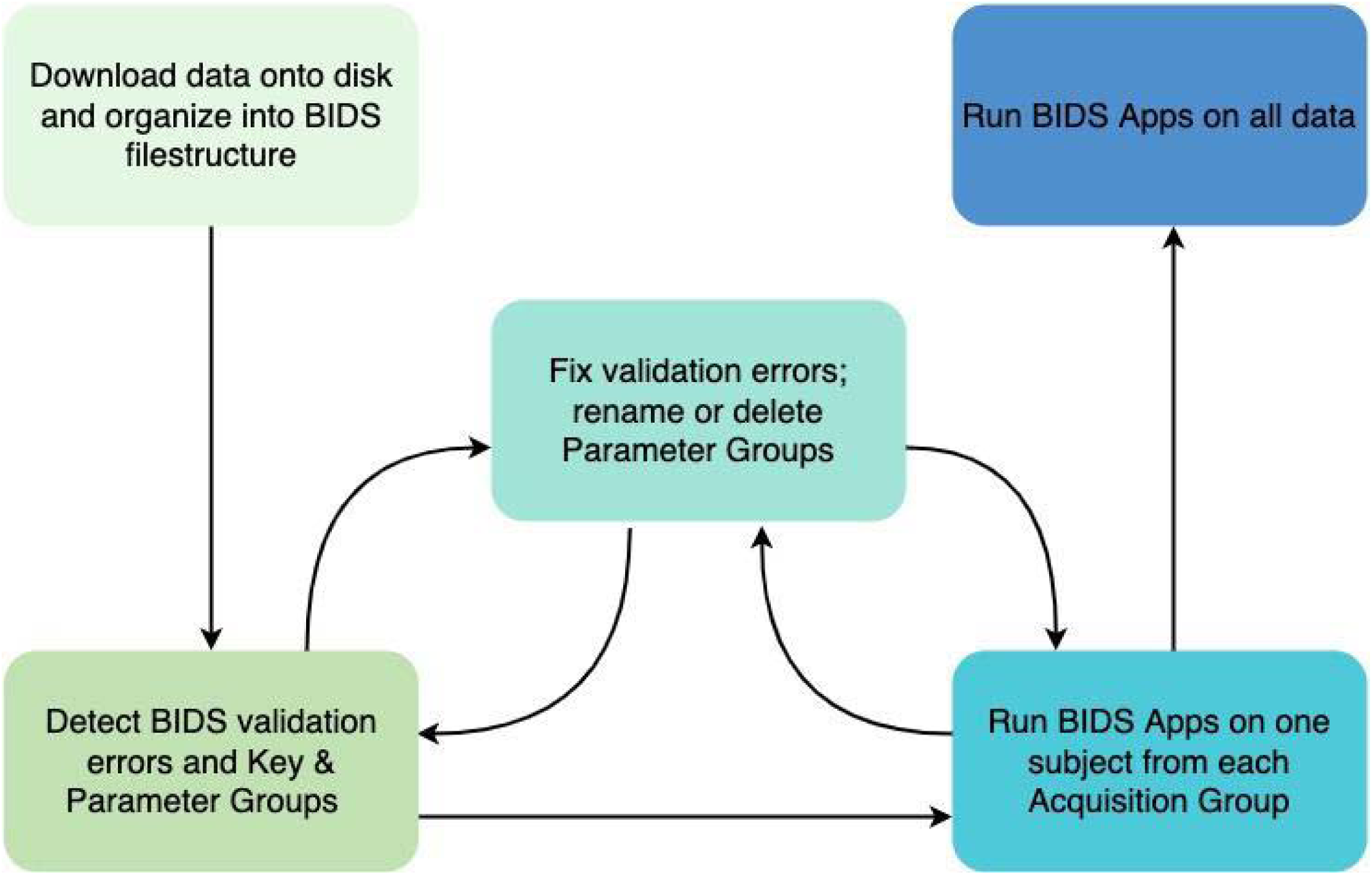
CuBIDS workflow. The CuBIDS workflow begins after the generation of NIfTI files and JSON sidecars and ends directly before the execution of pre-processing pipelines. We start with a BIDS dataset, which can be validated using CuBIDS’ robust version of the BIDS-validator. After purging the dataset of any sensitive fields, users can move to the next workflow stage: detecting Parameter Groups. Users can then rename or delete Parameter Groups. At any point in the workflow, users can implement version control to track changes made to the data using an easy-to-use, wrapped version of DataLad. Finally, users test one Exemplar Subject from each Acquisition Group on BIDS Apps to ensure each set of scanning parameters can run through pipelines error-free. CuBIDS includes command line programs for each step of the workflow (see **Table 1**).

### Accounting for NIfTI header information

Information from NIfTI headers—including number of volumes, voxel size, image dimensions, and image obliquity—is often important but is usually absent from JSON sidecars. We created a program, `cubids-add-nifti-info`, that reads information from the NIfTI header and adds it to the JSON sidecar. For example, knowing the number of volumes in a scan may be particularly useful when performing an initial quality assessment—i.e., identifying and removing scans with unexpectedly short durations (i.e., 20 volumes in an fMRI timeseries). Similarly, scans with vastly different voxel sizes or fields of view may be easily identified and removed if desired.

### BIDS validation

An essential first stage of curation is validation: finding the errors present in a BIDS dataset. This step is usually accomplished using the BIDS Validator. However, while BIDS validation is essential to the curation process, the standalone BIDS Validator can exhibit unstable file I/O behavior when validating large datasets (n>100). As a result, it sometimes fails unpredictably. To combat this issue, `cubids-validate` checks the BIDS layout using a wrapped, stable, scalable version of the standard BIDS Validator. To ensure scalability, `cubids-validate` parallelizes validation across participants, validating each subject directory on its own and deferring the detection of parameters that may vary across subjects. Thereafter, `cubids-validate` aggregates all validation errors found across participants in an easy-to-read CSV (see **Fig. 2**). This table includes one row for each file that contains a BIDS validation error and displays that filename along with a description of the error (see **Fig. 2**).

**Fig 2.**
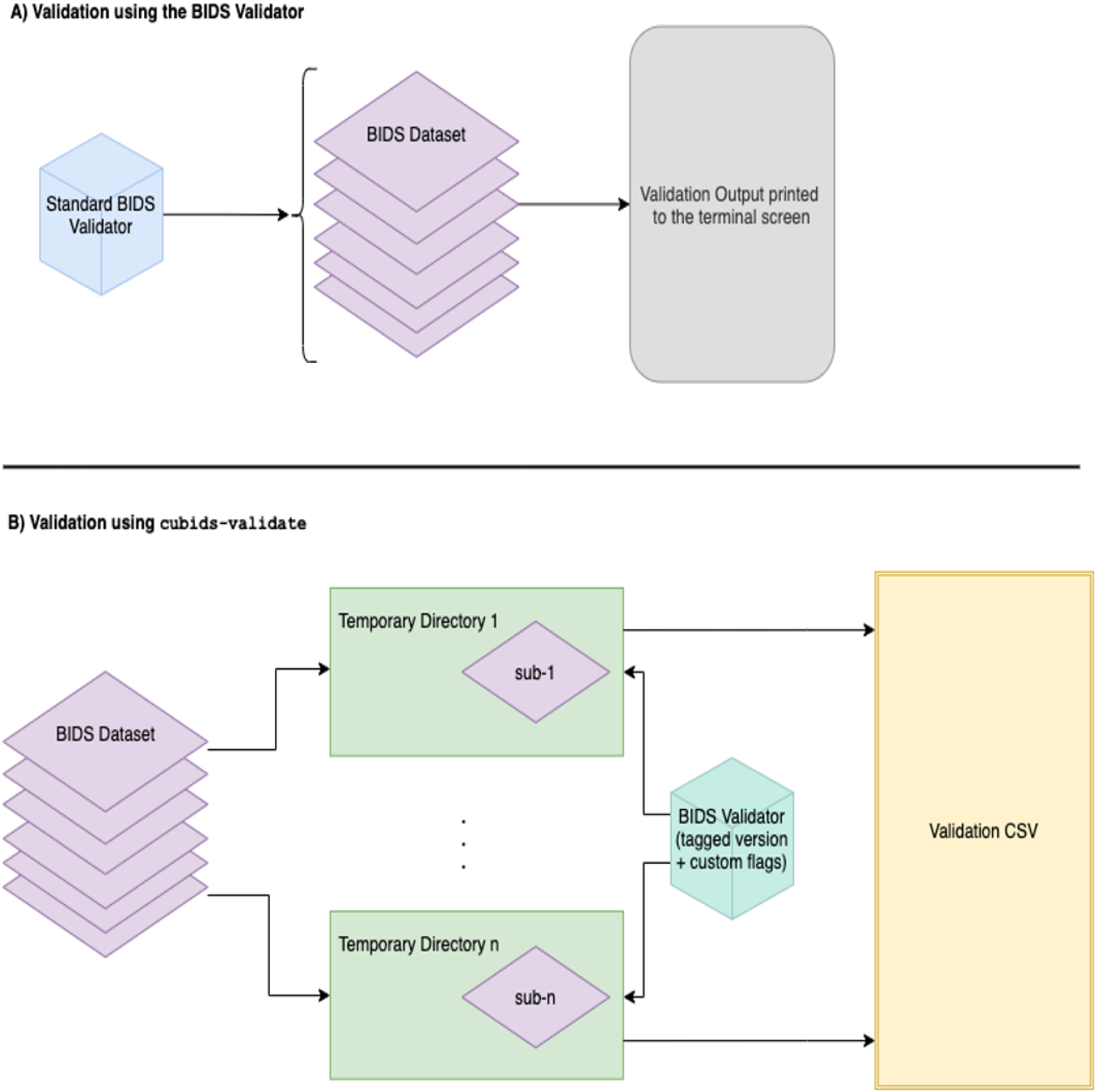
Stable, scalable BIDS validation. CuBIDS wraps a stable version of the BIDS Validator and adds a few additional features including the ability to reorganize the validator output into an easy to read, tabular structure and save it as a CSV. **A)** The standard BIDS Validator’ s default option (Gorgolewski et al., 2016) validates an entire BIDS dataset and outputs a summary of the errors and warnings it discovers to the terminal screen. **B)** In addition to visualizing the output in a scalable and easy-to-read format, `cubids-validate` includes the standard BIDS Validator’ s ability to ignore cross-session comparisons or metadata from NIfTI headers while also adding an option for sequential participant-by-participant validation. This feature, which we recommend users leverage, parallelizes validation and validates each subject directory as its own BIDS dataset.

In designing `cubids-validate`, we also intended to separate metadata heterogeneity detection from BIDS error detection. By default, the validator does both—providing large amounts of unactionable information concerning the metadata variance in the terminal output. For example, if a sample includes participants with different sets of scans, the standalone BIDS Validator will print warnings alerting the user to the presence of incongruencies across subjects, often producing copious output that can obscure critical issues. If there are errors or forms of inconsistency users would prefer excluded from the CuBIDS validation CSV, they can run `cubids-validate` with optional BIDS Validator flags such as `–-ignore_nifti_headers`, which disregards NIfTI header content during validation and `-- ignore_subject_consistency`, which we set as the default and skips checking that any given file for one subject is present for all other subjects. Furthermore, we implemented `-- sequential`, which parallelizes validation by running the BIDS Validator sequentially on each subject (see **Fig. 2B**), and `–-sequential-subjects`, which filters the sequential run to only include the listed subjects, e.g. `--sequential-subjects sub-01 sub-05 sub-09`. These flags allow users to focus the validation process exclusively on the issues and subjects they would like to evaluate, and the sequential option, which parallelizes validation, addresses the standalone BIDS Validator’ s scalability issue.

### Grouping: heterogeneity detection and classification

While `cubids-validate` will find and display BIDS validation errors present in a dataset, it does not identify metadata parameters that might be inconsistent or omitted. For this reason, we developed `cubids-group`: a grouping function that classifies the heterogeneity present in a BIDS dataset and displays it in readable CSVs. The input to `cubids-group` is the path to the root of a BIDS Dataset, and the program produces four outputs, each of which gives a different view of the underlying data. The first (and most important) is `summary.csv,` which contains one row per Parameter Group, and one column per metadata parameter present in the dataset. To understand the relative prevalence of each group, the program also counts, and includes in `summary.csv`, the number of files in each Key and Parameter Group; this documentation is very useful for visualizing metadata heterogeneity across the entire dataset.

The next output of `cubids-group` is `files.csv`, which contains one row per NIfTI file in the BIDS directory. This table keeps track of every scan’ s assignment to Key and Parameter Groups and includes a field that allows users to easily identify the Key and Parameter Groups to which each scan belongs. The next two grouping outputs organize the dataset by Acquisition Group. `AcqGrouping.csv` organizes the dataset by session and tags each one with its Acquisition Group number. Finally, `AcqGroupInfo.txt` lists all Key Groups that belong to a given Acquisition Group along with the number of sessions each group possesses.

When applied to large datasets, `cubids-group` will often reveal issues within a BIDS dataset that validation alone does not always catch. Such issues include missing sidecars, missing metadata parameters, and scans with low numbers of volumes or unusual image and voxel dimensions. For this reason, `cubids-group` can aid users in performing first pass quality assurance on their BIDS dataset. Since `summary.csv` breaks down the dataset by Parameter Group with one column per scanning parameter, users can then search that CSV by desired parameters. Next, users can set a threshold or requirement for a certain parameter (e.g. number of volumes or dimension/voxel size) and use `cubids-purge` to remove scans that do not possess the desired values for those parameters. For example, a user may want to remove all fMRI scans with a low number of volumes before data processing with a BIDS App such as fMRIPrep.

### Applying changes

The `cubids-apply` program provides an easy way for users to manipulate their datasets. Specifically, `cubids-apply` can rename files according to the users’ specification in a tracked and organized way. Here, the `summary.csv` functions as an interface modifications; users can mark Parameter Groups they want to rename (or delete) in a dedicated column of the `summary.csv` and pass that edited CSV as an argument to `cubids-apply.`

Additionally, `cubids-apply` can automatically rename files in Variant Groups based on their scanning parameters that vary from those in their Key Groups’ Dominant Parameter Groups. Renaming is automatically suggested when the `summary.csv` is generated from a `cubids-group` run, with the suggested new name listed in the CSV’ s “Rename Key Group” column. CuBIDS populates this column for all Variant Groups—e.g., every Parameter Group except the Dominant one. Specifically, CuBIDS will suggest renaming all Non-Dominant Parameter Groups to include VARIANT* in their acquisition field where * is the reason the Parameter Group varies from the Dominant Group. For example, when CuBIDS encounters a Parameter Group with a repetition time that varies from the one present in the Dominant Group, it will automatically suggest renaming all scans in that Variant Group to include `acquisition-VARIANTRepetitionTime` in their filenames. When the user runs `cubids-apply`, filenames will get renamed according to the auto-generated names in the “Rename Key Group” column in the `summary.csv` (see **Fig. 3**).

**Fig. 3.**
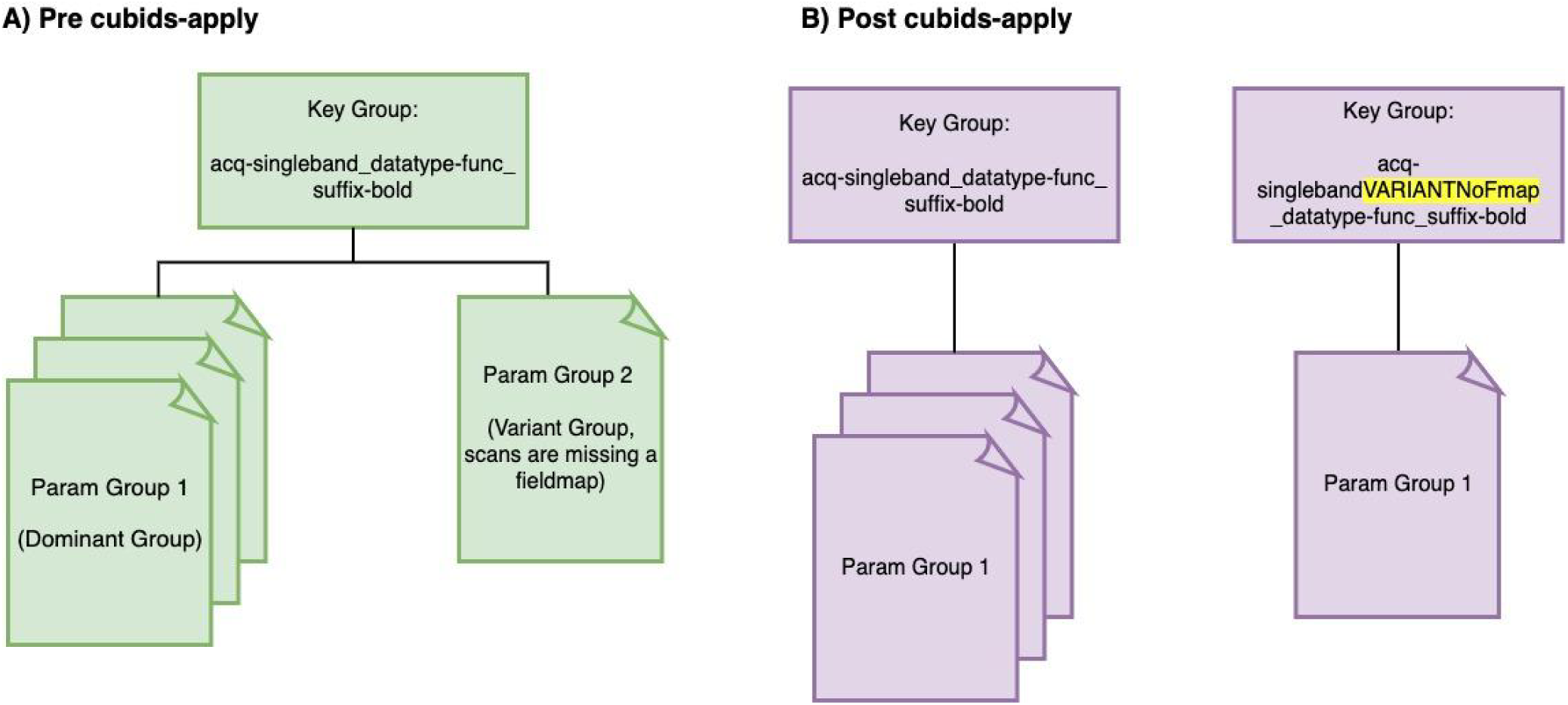
Parsing the dataset by Parameter Group. The `summary.csv` file is a `cubids-group` output that contains one row per Parameter Group and one column per scanning parameter. Thus, this CSV summarizes all metadata present within a dataset. **A)** Before `cubids-apply` is run, a given Key Group may have multiple Parameter Groups, each containing a different set of scanning parameters. This summary table includes a “Rename Key Group” column that auto-configures when `cubids-apply` is run and labels each non-dominant Parameter Group as a Variant Group based on the scanning parameters that differentiate that group from the Dominant Group. Specifically, CuBIDS represents this variance by adding “VARIANT*”––where * indicates the metadata parameters that cause those files to vary from the Dominant Group––to the “acq” field of those files in non-dominant Parameter Groups. For example, in **A)**, the metadata in the Param Group 2 scan differs from that of the Dominant Group (Param Group 1) scan because that scan is missing a fieldmap. The result of running `cubids-apply` can be seen in **B)** where the Param Group 2 scan ends up in a new Key group because CuBIDS added “VARIANTNoFmap” to the acquisition field of its filename when `cubids-apply` was run.

### Customizable configuration

CuBIDS also features an optional, customizable, modality-specific configuration file. This file can be passed as an argument to `cubids-group` and `cubids-apply` using the `–-config` flag and allows users to customize grouping settings based on modality and parameter. Each Key Group is associated with one (and only one) modality, as BIDS filenames include modality-specific values as their suffixes. This easy-to-modify configuration file provides several benefits to curation. First, it allows users to add and remove metadata parameters from the set that determines groupings. This can be very useful if a user deems a specific metadata parameter irrelevant and wishes to collapse variation based on that parameter into a single Parameter Group. Second, the configuration file allows users to apply tolerances for parameters with numerical values. This functionality allows users to avoid very small differences in scanning parameters (i.e., a TR of 3.0s vs 3.0001s) being split into different Parameter Groups. Third, the configuration file allows users to determine which scanning parameters are listed in the `acquisition` field when auto-renaming is applied to Variant Groups.

### Exemplar testing

In addition to facilitating curation of large, heterogeneous BIDS datasets, CuBIDS also prepares datasets for testing BIDS Apps. This portion of the CuBIDS workflow relies on the concept of the Acquisition Group: a set of sessions that have identical scan types and metadata across all imaging modalities present in the session set. Specifically, `cubids-copy-exemplars` copies one subject from each Acquisition Group into a separate directory, which we call an Exemplar Dataset. Since the Exemplar Dataset contains one randomly selected subject from each unique Acquisition Group in the dataset, it will be a valid BIDS dataset that spans the entire metadata parameter space of the full study. If users run `copy-exemplars` with the `–-use-datalad` flag, the program will ensure that the Exemplar Dataset is tracked and saved in DataLad. If the user chooses to forgo this flag, the Exemplar Dataset will be a standard directory located on the filesystem. Once the Exemplar Dataset has been created, a user can test it with a BIDS App (e.g., fMRIPrep or QSIPrep) to ensure that each unique set of scanning parameters will pass through the pipelines successfully. Because BIDS Apps auto-configure workflows based on the metadata encountered, they will process all scans in each Acquisition Group in the same way. By first verifying that BIDS Apps perform as intended on the small sub-sample of participants present in the Exemplar Dataset (that spans the full variation of the metadata), users can confidently move forward processing the data of the complete BIDS dataset.

## RESULTS

The CuBIDS workflow is currently being used in neuroimaging labs at a number of institutions including the University of Pennsylvania, the Children’ s Hospital of Philadelphia, the Child Mind Institute, and the University of Minnesota’ s Masonic Institute for the Developing Brain. To demonstrate the utility of CuBIDS, here we apply the software to two datasets. First, we curate a small example dataset that is included in the software’ s GitHub repository and can be downloaded <u>here</u>. Second, we apply CuBIDS to the large-scale data of the Philadelphia Neurodevelopmental Cohort.

### The CuBIDS workflow for curating a BIDS dataset (example dataset)

The following walkthrough displays the process of curating a dataset using CuBIDS on a Linux machine. This example walkthrough is also documented on the <u>CuBIDS Read the Docs page</u>. To do so, we use an example dataset that is bundled with the software. For this demonstration, we install CuBIDS inside a <u>conda environment</u>. Note that if you are using an Apple M1 chip machine, you will need to install <u>Miniforge</u> instead of Miniconda. Once we have conda installed we create and activate a new environment using the following commands:

~~~
conda create -n test-env python=3.8
conda activate test-env
~~~

To obtain CuBIDS locally, we can use pip to download <u>our software</u> from the Python Package Manager (Pypi) using the following command:

~~~
pip install CuBIDS
~~~

Alternatively, we can clone from the <u>CuBIDS GitHub repository</u> using the following command:

~~~
git clone https://github.com/PennLINC/CuBIDS.git
~~~

Now that we have a copy of the source code, we can install it by running

~~~
cd CuBIDS
pip install -e .
~~~

We will now need to install some dependencies of CuBIDS. To do this, we first must install <u>nodejs</u>. We can accomplish this using the following command:

~~~
conda install nodejs
~~~

Now that we have npm installed, we can install the bids-validator using the following command:

~~~
npm install -g bids-validator@1.7.2
~~~

In this example, we use the bids-validator v1.7.2. using a different version of the validator may result in slightly different validation csv printouts, but CuBIDS is compatible with all versions of the validator at or above v1.6.2. Throughout this example walkthrough, we use DataLad for version control, so we will also need to install both DataLad and git-annex, the large file storage software DataLad runs under the hood. Installation instructions for DataLad and git-annex can be found <u>here</u>.

Now that we have installed CuBIDS and all necessary dependencies, we are ready to begin the curation process on our example dataset. We create a `CuBIDS_Test` directory to function as the working directory and navigate to it as follows:

~~~
mkdir $PWD/CuBIDS_Test
cd CuBIDS_Test
~~~

Throughout this walkthrough, we will run all commands from the `CuBIDS_Test` directory. Next, we download `BIDS_Dataset.zip` (a ZipFile containing the example dataset) and unzip as follows:

~~~
curl –sSLO
https://github.com/PennLINC/CuBIDS/raw/main/cubids/testdata/BIDS_Dataset.zip
unzip BIDS_Dataset.zip
rm BIDS_Dataset.zip
~~~

As a first step, we use CuBIDS to identify the metadata fields present in the dataset. This is accomplished with the following command:

~~~
cubids-print-metadata-fields BIDS_Dataset
~~~

This command returns a total of 66 fields, including acquisition parameters and other metadata fields present in the dataset’ s JSON sidecars. Some of these fields contain simulated protected health information (PHI) such as `PatientName` that we wish to remove. Completing this step prior to checking the BIDS dataset into DataLad is critical, as we must ensure PHI is not tracked as part of version control. To remove the `PatientName` field from the sidecars, we can use the command:

~~~
cubids-remove-metadata-fields BIDS_Dataset --fields PatientName
~~~

If we were to run `cubids-print-metadata-fields` once more, we would see that `PatientName` is no longer present in the dataset. Now that all PHI has been removed from the metadata, we are ready to check our dataset into DataLad. To do this, we run the following command:

~~~
datalad create -c text2git BIDS_Dataset_DataLad
~~~

The creation of our DataLad dataset will be accordingly reflected in the dataset’ s version control history, or “git log” (see example in **Fig. 4A**). At any point in the CuBIDS workflow, we can view a summary of our dataset’ s version history by running the following commands:

**Fig. 4.**
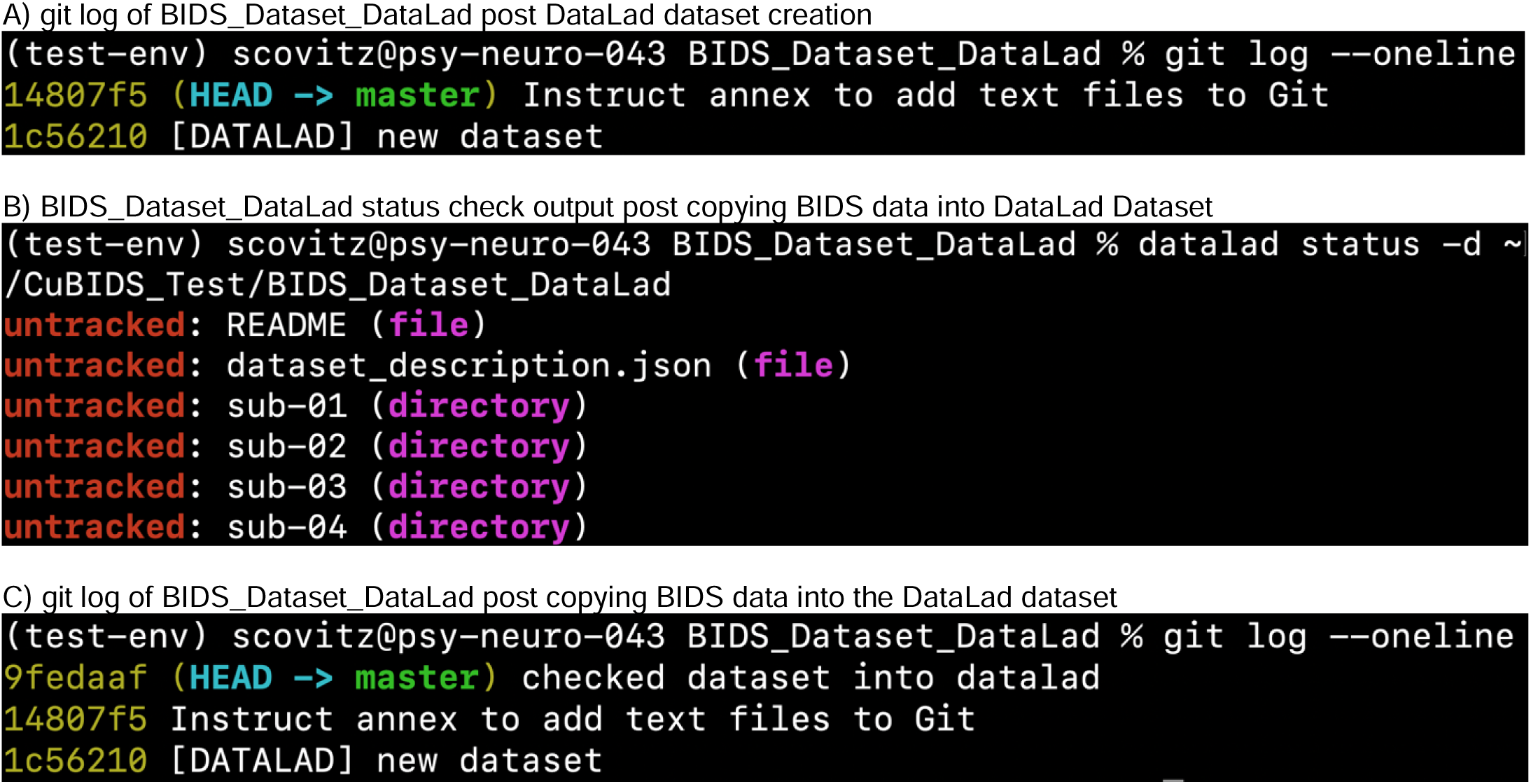
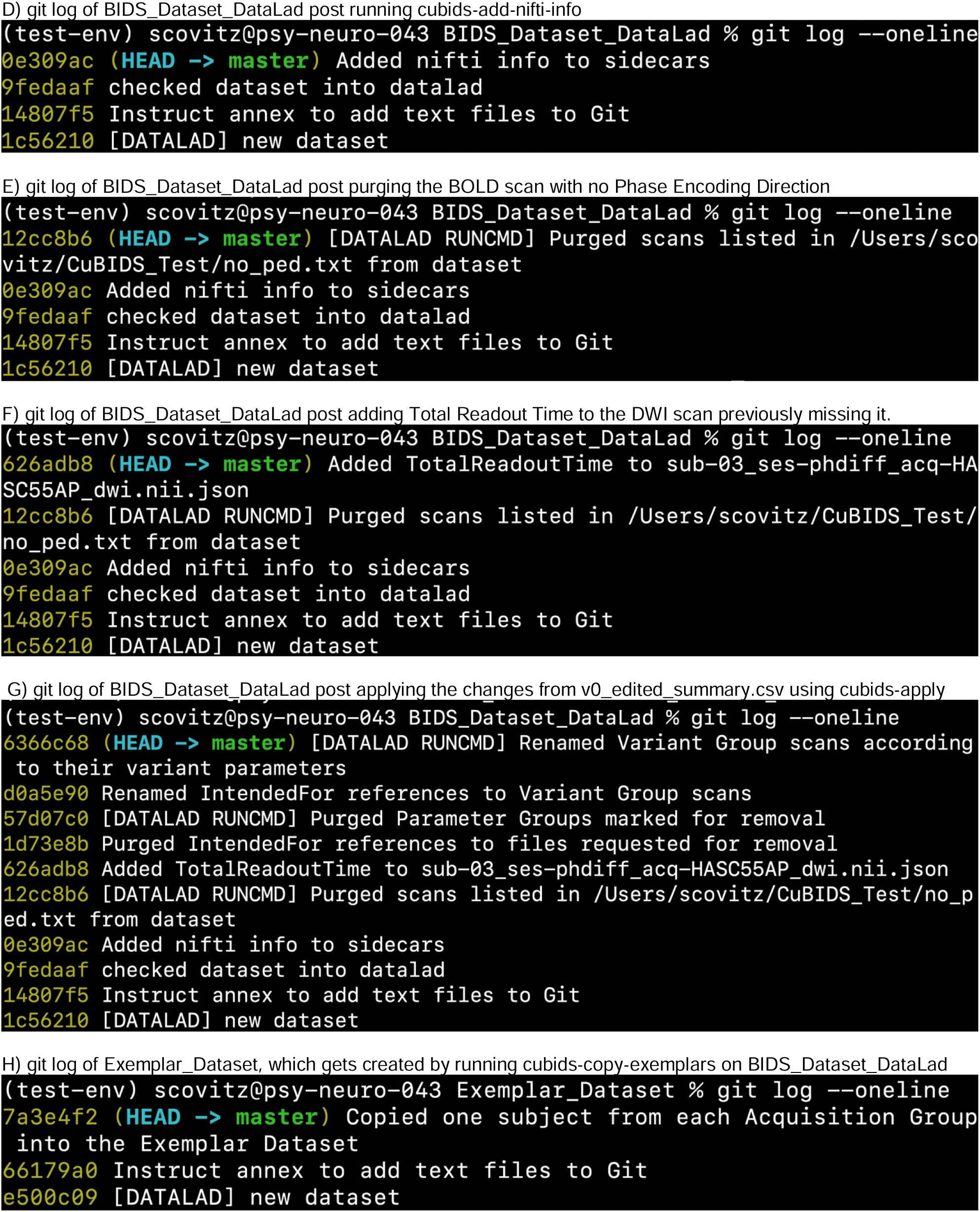
Version history throughout the curation process. The above panels display the version history of the small, example DataLad dataset we curated to display the effectiveness of the CuBIDS workflow. These panels are screenshots of the git history of the dataset taken after each change was made to the data. A shasum (yellow string of letters and numbers to the left of each commit message) is assigned to each commit, and each commit is recorded with a message (white text describing the changes made to the data). If users would like more information about each commit, they can run the `git log` command without the `oneline` flag to get a detailed summary of each commit. This summary will include files that were changed, exact changes that were made to each file, date and time of the commit, and information about the git user who made the changes. At any point in the workflow after checking the dataset into DataLad, we can use `cubids-undo` to revert the dataset back to the previous commit.

~~~
cd BIDS_Dataset_DataLad
git log --oneline
cd ..
~~~

Next, we copy the contents of our BIDS dataset into the newly created and currently empty DataLad dataset:

~~~
cp -r BIDS_Dataset/* BIDS_Dataset_DataLad
~~~

In addition to being able to access the version history of our data, any point in this workflow, we can also check the status of untracked (not yet saved) changes using the `datalad status` command, as seen below:

~~~
datalad status -d BIDS_Dataset_DataLad
~~~

This command produces a description of the changes we have made to the data since the last commit (see **Fig. 4B**). The command above shows all files untracked, as we have copied the BIDS data into `BIDS_Dataset_DataLad` but have not yet saved those changes. Our next step is to run save. It is best practice to provide a detailed commit message, for example:

~~~
datalad save -d BIDS_Dataset_DataLad -m “checked dataset into datalad”
~~~

This commit is reflected in our git log (see **Fig. 4C**). Now that the dataset is checked into DataLad, at any point in the workflow going forward, we can run the following command to revert the dataset back to the previous commit:

~~~
cubids-undo BIDS_Datast_DataLad
~~~

At this stage, we also recommend removing the ‘‘BIDS_Dataset’’ directory — its contents are safely copied into and tracked in ‘‘BIDS_Dataset_DataLad’’.

Next, we seek to add new fields regarding our image parameters that are only reflected in the NIfTI header to our metadata; these include important details such as image dimensions, number of volumes, image obliquity, and voxel sizes. To do this, we run:

~~~
cubids-add-nifti-info BIDS_Dataset_DataLad --use-datalad
~~~

This command adds the NIfTI header information to the JSON sidecars and saves those changes. In order to ensure that this command has been executed properly, we can run `cubids-print-metadata-fields` once more, which reveals that NIfTI header information has been successfully included in the metadata. Since we ran `add-nifti-info` with the `use-datalad` flag set, CuBIDS will automatically save the changes made to the dataset to the git log (see **Fig. 4D**).

The next step in the CuBIDS workflow is to understand what BIDS validation errors may be present (using `cubids-validate`) as well as the structure, heterogeneity, and metadata errors present in the dataset (using `cubids-group`). Notably, neither of these two programs requires write access to the data, as each simply reads in the contents of the data and creates CSVs that parse the metadata and validation errors present. Validation can be accomplished by running the following command:

~~~
cubids-validate BIDS_Dataset_DataLad v0 –sequential
~~~

The use of the `sequential` flag forces the validator to treat each participant as its own BIDS dataset. This command produces `v0_validation.csv` (see **Supplementary Data 1A)**.

This initial validation run reveals that Phase Encoding Direction (PED) is not specified for one of the BOLD task-rest scans. We can clearly see that we either need to find the PED for this scan elsewhere and edit that sidecar to include it or remove that scan from the dataset, as this missing scanning parameter will render field map correction impossible. For the purpose of this demonstration, we elect to remove the scan. To do this, we run the following command:

~~~
cubids-purge BIDS_Dataset_DataLad no_ped.txt --use-datalad
~~~

Here, `no_ped.txt` (see **Supplementary Data 1B**) is a text file containing the **full path** to the dwi scan flagged in `v0_validation.txt` for missing PED. The user must create this file before running `cubids-purge` (a command such as `echo $PWD/BIDS_Dataset_DataLad/sub-02/ses-phdiff/func/sub-02_ses-phdiff_task-rest_bold.nii.gz > no_ped.txt` will work).

We elect to use `cubids-purge` instead of simply removing the scan because `cubids-purge` will ensure all associated files, including sidecars and IntendedFor references in the sidecars of fieldmaps, are also deleted. This change will be reflected in the git history (see **Fig. 4E)**.

Returning again to `v0_validation.csv`, we can also see that there is one DWI scan missing TotalReadoutTime, a metadata field necessary for certain pipelines. In this case, we determine that TotalReadoutTime (TRT) was erroneously omitted from the DWI sidecars. For the purpose of this example, we assume we are able to obtain the TRT value for this scan (perhaps by asking the scanner technician). Once we have this value, we manually add it to the sidecar for which it is missing by opening `BIDS_Dataset_DataLad/sub-03/ ses-phdiff/dwi/sub-03_ses-phdiff_acq-HASC55AP_dwi.json` in an editor and adding the following line:

~~~
“TotalReadoutTime”: 0.0717598,
~~~

on a new line anywhere inside the curly braces between lines containing parameters and their values, save the changes, and close the JSON file. We then save the latest changes to the dataset with a detailed commit message as follows:

~~~
datalad save -d BIDS_Dataset_DataLad -m “Added TotalReadoutTime to sub-03_ses-phdiff_acq-HASC55AP_dwi.nii.json”
~~~

This change will be reflected in the git history (see **Fig. 4F**).

To verify that there are no remaining validation errors, we rerun validation with the following command:

~~~
cubids-validate BIDS_Dataset_DataLad v1 –sequential
~~~

This command will produce no CSV output and instead print `“No issues/warnings parsed, your dataset is BIDS valid”` to the terminal, which indicates that the dataset is now free from BIDS validation errors and warnings.

Along with parsing the BIDS validation errors in our dataset, it is important to understand the dataset’ s structure, heterogeneity, and metadata errors. To accomplish these tasks, we use `cubids-group`. Large datasets almost inevitably contain multiple validation and metadata errors. As such, it is typically useful to run both `cubids-validate` and `cubids-group` in parallel, as validation errors are better understood within the context of a dataset’ s heterogeneity. Additionally, being able to see both the metadata errors that grouping reveals alongside BIDS errors that the validator catches gives users a more comprehensive view of the issues they will need to fix during the curation process. Note that `cubids-group` requires <u>full paths</u> to both the BIDS Dataset and the output prefix. The command to run the grouping function is as follows:

~~~
cubids-group $PWD/BIDS_Dataset_DataLad $PWD/v0
~~~

As noted in **Table 1**, this command will produce four tables that display the dataset’ s heterogeneity in different ways. First, `v0_summary.csv` contains all detected Key and Parameter groups and provides a high-level overview of the heterogeneity in the entire dataset (see **Supplementary Data 1C**). Second, `v0_files.csv` (see **Supplementary Data 1D**) maps each imaging file in the BIDS directory to a Key and Parameter group. Third, `v0_AcqGrouping.csv` (see **Supplementary Data 1E**) maps each session in the dataset to an Acquisition Group. Finally, `v0_AcqGroupInfo.txt` (see **Supplementary Data 1F**) lists the set of scanning parameters present in each Acquisition Group.

The next step in the CuBIDS curation process is to examine `v0_summary.csv`, which allows for automated metadata quality assurance (QA)––the identification of incomplete, incorrect, or unusable parameter groups based on acquisition fields such as dimension and voxel sizes, number of volumes, etc. While `v0_validation.csv` identified all BIDS validation errors present in the dataset, it will not identify several issues that might be present with the sidecars. Such issues include instances of erroneous metadata and missing sidecar fields, which may impact successful execution of BIDS Apps.

Examining `v0_summary.csv` (see **Supplementary Data 1C**) we can see that one DWI Parameter Group––`acquisition-HASC55AP_datatype-dwi_suffix-dwi 2`– –contains only one scan (see “Counts” column) with only 10 volumes (see “NumVolumes” column). Since the majority of DWI scans in this dataset have 61 volumes, CuBIDS assigns this single scan to a “Non-Dominant”, or “Variant” Parameter Group and populates that Parameter Group’ s “RenameKeyGroup” column in `v0_summary.csv` with `acquisition-HASC55APVARIANTNumVolumes_datatype-dwi_suffix-dwi`. For the purpose of this demonstration, we elect to remove this scan because it does not have enough volumes to be usable for most analyses. To do this, we can either use `cubids-purge`, or we can edit `v0_summary.csv` by adding “0” to the “MergeInto” column in the row (Parameter Group) we want to remove. For this walkthrough, we chose the latter. To do this, we open `v0_summary.csv` in an editor, navigate to row 4, which contains all information for Key Group `acquisition-HASC55AP_datatype-dwi_suffix-dwi` Parameter Group 2. If we scroll to the `NumVolumes` column (row 4, column S), we see this Parameter Group has only 10 volumes, which explains why it received an auto-generated Rename Key Group value of `acquisition-HASC55APVARIANTNumVolumes_datatype-dwi_suffix-dwi`. Remaining in this same row, we navigate back to column C, which is labeled “MergeInto” and manually a “0” to the cell in row 4 column C. This will ensure all scans in that Parameter Group (in this example, just one scan) are removed when we run `cubids-apply`. We then export and save the CSV in our `CuBIDS_Test` working directory as `v0_edited_summary.csv` (see **Supplementary Data 1G**). We will then save this edited version of `v0_summary.csv` as `v0_edited_summary.csv`, which will be passed into `cubids-apply` in our next curation step.

Now that all metadata issues have been remedied––both the validation and summary outputs appear problem-free––we are ready to rename our files based on their Rename Key Group values and apply the requested deletion in `v0_edited_summary.csv`. The `cubids-apply` function renames scans in each Variant Parameter Group according to the metadata parameters with a flag “`VARIANT`”, which is useful because the user will then be able to see, in each scan’ s filename, which metadata parameters associated with that scan vary from those in the acquisition’ s Dominant Group. Note that like in `cubids-group`, `cubids-apply` requires <u>full paths</u> to the BIDS Dataset, summary and files CSVs, and output prefix. We execute `cubids-apply` with the following command:

~~~
cubids-apply $PWD/BIDS_Dataset_DataLad
$PWD/v0_edited_summary.csv $PWD/v0_files.csv $PWD/v1 --use- Datalad
~~~

Checking our git log, we can see that all changes from apply have been saved (see **Fig. 4G**). As a final step, we can check the four grouping CSVs `cubids-apply` produces (see **Supplementary Data 1H-K**) to ensure they look as expected––that all files with variant scanning parameters have been renamed to indicate the parameters that vary in the acquisition fields of their filenames (and therefore Key Group names).

At this stage, the curation of the dataset is complete; next is pre-processing. CuBIDS facilitates this subsequent step through the creation of an Exemplar Dataset: a subset of the full dataset that spans the full variation of acquisitions and parameters by including one subject from each Acquisition Group. By testing only one subject per Acquisition Group, users are able to pinpoint both the specific metadata values and scans that may be associated with pipeline failures; these acquisition groups could then be evaluated in more detail and flagged for remediation or exclusion. The Exemplar Dataset can easily be created with the `cubids-copy-exemplars` command, to which we pass in `v1_AcqGrouping.csv`––the post-apply acquisition grouping CSV (see **Supplementary Data 1J**).

~~~
cubids-copy-exemplars BIDS_Dataset_DataLad Exemplar_Dataset v1_AcqGrouping.csv --use-datalad
~~~

Since we used the `use-datalad` flag, `Exemplar_Dataset` is a DataLad dataset with the version history tracked in its git log (see **Fig. 4H**). Once a preprocessing pipeline completes successfully on the Exemplar Dataset, the full dataset can be executed with confidence, as a pipeline’ s behavior on the full range of metadata heterogeneity in the dataset will have already been discovered during exemplar testing.

### Application to a large-scale study of brain development

In addition to applying the CuBIDS workflow to a toy dataset, here we describe the workflow applied to the Philadelphia Neurodevelopmental Cohort (PNC), a multimodal dataset of n=1,601 participants. This set of steps involved iterative rounds of checking and fixing due to the heterogeneity and size of the dataset, so the following section will be a summary of how we used CuBIDS to curate this dataset (rather than a step-by-step walkthrough).

Examining the initial summary table (see **Supplementary Data 2B**), we find that PNC contains 144 Parameter Groups––scans containing both identical BIDS filename key-value pairs and identical metadata parameters present in their sidecars. The summary table is organized by modality, so we can easily see that some modalities and acquisitions in the dataset are much more heterogeneous with respect to their metadata parameters than are others. For example, from the table, we can see that PNC has only one Key Group for T1w scans and only three Parameter Groups. Furthermore, according to the “Counts” column of the csv, the vast majority of T1w scans (n=1597), are in the Dominant Parameter Group. By contrast, in this same summary table we can see that there are 62 different Parameter Groups in the dataset for task-frac2back BOLD fMRI scans. Examining the “RenameKeyGroup” column of those frac2back rows in the summary table, we can see that the primary source of variance is the number of volumes acquired.

Our team relied upon the summary table to make a number of curation decisions–– especially inclusion/exclusion based on metadata. The summary table provided a platform for collaboration and discussion among the team that was curating, validating, and modifying the dataset. Since PNC was curated with DataLad and is saved as a DataLad Dataset, a detailed history of the curation decisions can be found in the commit history. Each change to the dataset was saved with a commit message, so all modifications we made to the dataset are tracked and tagged with a shasum. If we want to restore PNC to a previous curation stage, we can so using `cubids-undo`.

Modifications to PNC during the curation stage, documented in the dataset’ s git log, include creating and adding previously missing events tsvs, adding NIfTI header information to all sidecars in the dataset (using the `cubids-add-nifti-info` command), removing DWI scans that do not have enough volumes to successfully run through diffusion preprocessing pipelines (e.g. QSIPrep), and adding Parallel Reduction Factor in Plane to two sidecars that were initially missing this field. Once the validation and grouping outputs revealed we had no more BIDS and metadata issues to fix, we ran `cubids-apply` on the dataset to produce a new set of CSVs (see **Supplementary Data 2G-J**). As can be seen in the summary table (see **Supplementary Data 2G**) generated by `cubids-apply`, all scans in Variant (e.g., non-Dominant) Groups were renamed based on the scanning parameters that are variant (see **Fig. 3**).

We then executed the final step of the CuBIDS workflow, which entailed running `cubids-copy-exemplars`. This command creates the Exemplar Dataset––a DataLad-tracked BIDS dataset containing one subject from each Acquisition Group (see **Supplementary Data 2I**). The final, curated version of PNC contains 1601 participants, 15,077 scans, and 65 Acquisition Groups. Thus, the Exemplar Dataset contains only 65 participants but spans the entire dataset’ s parameter space, reducing the scope of the pipeline testing by 96%. We then used this Exemplar Dataset to test modality-specific preprocessing pipelines such as fMRIPrep and QSIPrep.

For large datasets especially, exemplar testing can be a necessary step; users will often need to go back and re-curate aspects of the BIDS data based on metadata errors that only become apparent during pipeline runs on the Exemplar Dataset. For example, after we ran the PNC Exemplar Dataset through QSIPrep, we noticed that one participant (whose Acquisition Group includes 34 participants), failed to complete the pipeline successfully. After examining the error log, we realized that for this participant, the number of bvals did not match the number of volumes in the scan’ s sidecar (which was pulled directly from the NIfTI header and added to the sidecar during the `cubids-add-nifti-info` step of curation). Since all participants in the same Acquisition Group possess identical scanning parameters, when a pipeline encounters a metadata error in an Exemplar Subject, all participants in that Acquisition Group will have that same error and thus require the same fix. Accordingly, we used `cubids-purge` to remove all DWI scans from that Exemplar Group and reran `cubids-group` to obtain our final CuBIDS outputs (see **Supplementary Data 2K-N**). Since all other Exemplar Subjects passed through QSIPrep successfully, we were then able to run the pipeline through the entire dataset without concern that erroneous metadata would impact preprocessing.

## DISCUSSION

Ample recent evidence has emphasized the challenges to reproducibility in neuroimaging research (Kang et al., 2016). Although often overlooked, curation can be a critical part of the scientific workflow. Because curation is often the first step after data acquisition, errors in curation can ramify throughout each subsequent stage. BIDS apps adapt to metadata encountered in an automatic and flexible way, which can be a vulnerability in ensuring datasets are processed identically. If BIDS data are improperly curated, pre-processing pipelines may mis-configure, with the potential to impact eventual results. Curation challenges are particularly acute in large-scale data resources, which continue to proliferate (e.g., UK Biobank, ABCD, PNC, HBN, HCP, etc.) (Bycroft et al., 2018; Karcher et al., 2021; Satterthwaite et al., 2013; Alexander et al., 2017; Van Essen et al., 2013). In large datasets, curation is often an iterative, manual process that is neither well documented nor reproducible. To address these challenges, CuBIDS allows for reproducible data curation at scale. As discussed below, our software provides five main advantages: stability in validation, reproducibility in curation, the ability to identify and manage heterogeneity, transparency in naming, and accelerated pipeline testing.

### Stable BIDS validation at scale

The BIDS Validator is the current standard tool for validation of all BIDS datasets. It is widely used and plays an essential role in the standard BIDS workflow; it effectively identifies the ways in which a dataset does not comply with BIDS standards. However, it does not scale well, at times failing unpredictably on larger samples. Furthermore, when run in a Linux shell, the validator prints (often a large volume) of text describing the errors and warnings to the terminal screen. For large datasets with many errors and warnings, such information is often quite difficult to visualize and comprehend. We wrapped the BIDS Validator in the `cubids-validate` CLI to address these challenges, creating a scalable implementation that yields a readable CSV. This allows users to easily identify the range of validation issues that may be present in a large-scale dataset.

### Reproducible data curation

Curation of large, heterogeneous BIDS datasets is an iterative, multi-step task. However, this process is often not reproducible, which, in turn, may compromise the reproducibility of subsequent workflows. Without version control, any decision made during the curation process– –such as inclusion/exclusion decisions or editing metadata––will go unrecorded. Further, if the person curating the data makes a mistake, they will have no clear way to undo that mistake and revert the data to a prior state. In leveraging CuBIDS’ use of DataLad, users can save each change made to a dataset with a detailed commit message (e.g. “Removed all DWI scans with less than 30 volumes”). If a user erroneously changes the data and wants to undo those changes, `cubids-undo` reverts the most recent commit.

### Parsing heterogeneity in large-scale data resources

While `cubids-validate` will catch instances where the data does not comply with BIDS format, it has important limitations. For example, validation does not always account for missing JSON sidecars or empty NIfTI headers. In addition, it will neither identify scans that have errant metadata values nor those with parameters that might render the scans unusable. This functionality is provided by `cubids-group`, which produces parameter-based summary tables that parse the dataset based on metadata, allowing for users to visualize and assess metadata quality in ways that validation cannot.

This functionality is especially critical in the curation of large datasets. Scaling up both the number of participants and the number of scanners within a single data resource has the potential to introduce a massive amount of heterogeneity to that study’ s eventual BIDS dataset. Heterogeneity in scanning parameters can result in heterogeneity in preprocessing pipelines; if users are do not appreciate the metadata heterogeneity in their dataset, they may be surprised by inconsistencies in preprocessing settings and outcomes. Further, parameter groups could be explicitly modeled when accounting for batch effects rather than just using scanner or site. Thus, being able to identify and correct metadata errors in a heterogeneous dataset is a critical part of the data curation process, as such decisions may impact the derived images from preprocessing pipelines.

### Enhancing transparency with Dominant and Variant Groups

In order to provide transparent documentation of parameter heterogeneity, the `cubids-apply` function renames scans in each modality according to their variant metadata parameters. For example, if the majority of BOLD task-rest scans in a dataset are Oblique but sub-X’ s scan is not oblique, CuBIDS users can choose to accept and apply the automatically suggested renaming of “`acq-VARIANTObliquity`” to that scan’ s filename. When performing sensitivity analyses on derivatives from datasets that have been curated using CuBIDS, researchers may choose to exclude any scans in Non-Dominant Groups to ensure that scanning parameters variance does not affect their results. Alternatively, researchers could use image harmonization tools (e.g., ComBat) to ensure that such variation does not impact analyses (Fortin et al., 2018).

### Accelerating pipeline testing with exemplar datasets

Even after careful curation, the best way to verify successful image processing is empirical testing. In a highly heterogeneous dataset, pipeline testing often reveals errors that were not immediately apparent on initial curation, which usually require minor additional adjustments to the metadata or exclusion of specific scans. Finding such edge cases while processing a large dataset can slow down the workflow, so it is advantageous to conduct pipeline testing before full deployment on a large data set. CuBIDS facilitates this process through the creation of Exemplar Datasets that include data from each Acquisition Group and thus span the full variation of the metadata present. After successful testing on the Exemplar Dataset, the likelihood that unexpected outcomes occur when the full dataset is processed is dramatically reduced. Furthermore, resource usage can be monitored during the exemplar runs to estimate the runtime and storage demands for processing the entire dataset.

### Limitations

CuBIDS possesses several limitations that should be acknowledged. First, at present, CuBIDS does not possess a GUI, so running the software requires basic knowledge of the terminal and Linux machines. However, such skills are likely to be a prerequisite for curating large-scale imaging datasets. Second, if users are curating BIDS Datasets with n>2,500 participants and using the DataLad-enabled version control option, CuBIDS programs that rely on saving changes made to the dataset might experience runtimes that extend to over an hour–– due to the need for DataLad to index such a large dataset. Third, and finally, CuBIDS is currently only able to be run on disk––either a local machine or a high performance computing cluster; users cannot currently run CuBIDS using either cloud-based computing (e.g., Amazon’ s S3) or neuroimaging databases such as the eXtensible Neuroimaging Archive Toolkit (XNAT), Longitudinal Online Research and Imaging System (LORIS), Collaborative Informatics Neuroimaging Suite (COINS), and the commercial platform Flywheel (Herrick et al., 2016; Das et al., 2012; Landis et al., 2016). Such functionality may be added to the software in future releases.

## CONCLUSIONS

Curating large, heterogeneous neuroimaging datasets can be a difficult and frustrating task. As the size and heterogeneity of data resources continues to expand, tools that allow for reproducible curation are not only helpful but necessary. CuBIDS facilitates efficient identification and correction of issues present in the metadata of heterogeneous BIDS datasets in a fully reproducible manner. Furthermore, CuBIDS Exemplar Datasets allow users to verify that BIDS Apps perform as intended on a small sub-sample of participants that spans the entire parameter space of the dataset, accelerating the processing of all data from the complete study. Together, CuBIDS allows users to simultaneously streamline curation and ameliorate metadata issues while maximizing reproducibility.

## ACKNOWLEDGEMENTS

This study was supported by grants from the National Institutes of Health: R01MH120482, R37MH125829, R01MH113550, R01EB022573, RF1MH116920, R01MH112847. Additional support was provided by the the CHOP-Penn Lifespan Brain Institute, the Penn Brain Science Center, and the Center for Biomedical Image Computing and Analytics

## APPENDIX A. SUPPLEMENTARY MATERIAL

**Supplementary Data 1 (for toy dataset in Results)**

Supplementary Data files 1A-O are zipped into **Supplementary_Data_1.zip** (submitted as supplementary material)

**Supplementary Data 2 (for PNC curation in Results)**

Supplementary Data files 2A-N are zipped into **Supplementary_Data_2.zip** (submitted as supplementary material)

## Notes

https://cubids.readthedocs.io/en/latest/

https://github.com/PennLINC/CuBIDS

https://pypi.org/project/cubids/

### Competing Interest Statement

The authors have declared no competing interest.

